# Mapping caudolenticular gray matter bridges in the human brain striatum through diffusion magnetic resonance imaging and tractography

**DOI:** 10.1101/2023.08.28.555179

**Authors:** Graham Little, Charles Poirier, Martin Parent, Laurent Petit, Maxime Descoteaux

**Affiliations:** Department of Computer Science, Université de Sherbrooke, Sherbrooke, Quebec, Canada; Université Bordeaux, CNRS, CEA, IMN, GIN, UMR 5293, F-33000 Bordeaux, France; CERVO Brain Research Center and Department of Psychiatry and Neuroscience, Faculty of Medicine, Université Laval, Quebec City, QC, Canada

## Abstract

In primates, the putamen and the caudate nucleus are connected by ∼1mm-thick caudolenticular gray matter bridges (CLGBs) interspersed between the white matter bundles of the internal capsule. Little is understood about the functional or microstructural properties of the CLGBs. In studies proposing high resolution diffusion magnetic resolution imaging (dMRI) techniques, CLGBs have been qualitatively identified as an example of superior imaging quality, however, the microstructural properties of these structures have yet to be examined. In this study, it is demonstrated for the first time that dMRI is sensitive to an organized anisotropic signal oriented in the direction parallel to the CLGBs, suggesting that dMRI could be a useful imaging method for probing the microstructure of the CLGBs. To demonstrate the anisotropic diffusion signal is coherently organized along the extent of the CLGBs and to enable a subsequent CLGB microstructural measurement, a novel tractography algorithm is proposed that utilizes the shape of the human striatum (putamen + caudate nucleus) to reconstruct the CLGBs in 3D. The method is applied to two publicly available and one locally acquired diffusion imaging datasets varying in resolution and imaging quality, thereafter demonstrating that the reconstructed CLGBs directly overlap expected gray matter regions in the human brain. In addition, the method is shown to accurately reconstruct CLGBs repeatedly across multiple test-retest cohorts. The novel tractography CLGB reconstructions are then used to extract a quantitative measurement of microstructure from a local model of the diffusion signal along the CLGBs themselves. This is the first work to comprehensively study the CLGBs in-vivo using dMRI and presents techniques suitable for future human neuroscience studies targeting these structures.

## 1 Introduction

Receiving massive projections from the entire cortical mantle, the striatum is often considered as the main input structure of the basal ganglia. During development, an increasing number of white matter fibers arising from virtually all areas of the cerebral cortex travel through the striatum (“The Human Central Nervous System, 4th ed.,” 2008). In primates including humans, many of these fibers converge to form the internal capsule, dividing the striatum into a dorsomedial caudate nucleus and a ventro-lateral putamen. Interestingly, the caudate nucleus and the putamen remain connected along the antero-posterior axis where striatal cell bridges are interspersed between bundles of the internal capsule forming what are named the caudolenticular gray matter bridges (CLGBs) (Dang et al., 2023). Little is understood about the functional or microstructural properties of CLGBs, other than they might receive significant afferent projections from the pre-supplementary motor area of the cerebral cortex (Inase et al., 1999). For reference, the CLGBs are visualized as an artistic rendering and on a histological slice in Figure 1.

**Figure 1:**
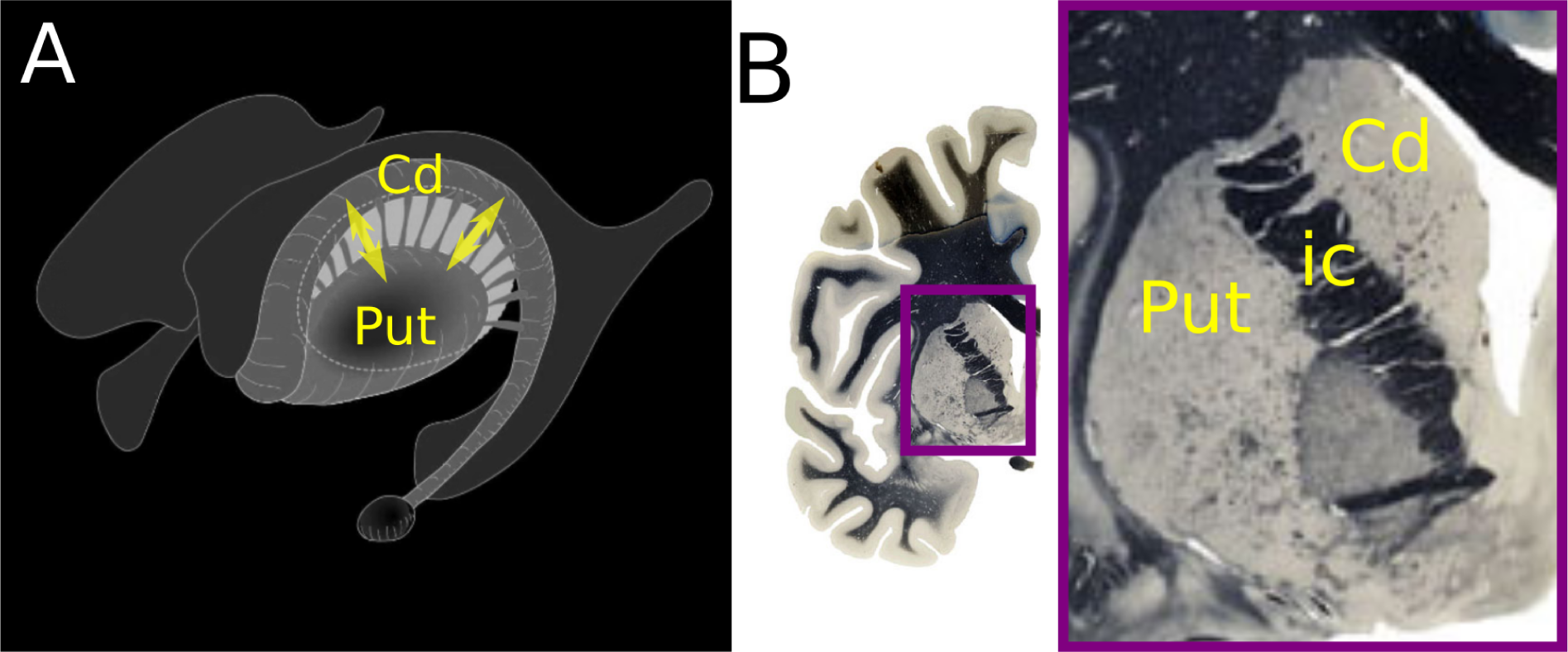
Visualization of the caudolenticular gray matter bridge (CLGB) anatomy. (A) Anatomically, the CLGBs (general orientation yellow arrows) are an extension of the gray matter of the striatum attaching the putamen (Put) and the caudate nucleus (Cd)(“Anatomical diagrams of the brain | e-Anatomy” https://doi.org/10.37019/e-anatomy/49401). (B) CLGBs crossing the internal capsule (ic) on a coronal histological section dissected from a single fixed brain from the Vogt brain collection (http://www.thehumanbrain.info/brain/sections.php).

Using magnetic resonance imaging (MRI), CLGBs can be visually depicted in the human brain at high resolutions (∼0.75 mm isotropic) on structural imaging (see example of T2 weighted image in Figure 2A) and parametric maps generated by diffusion tensor imaging (DTI) (Figure 2B). In this example, structural and diffusion imaging acquired in-vivo (Wang et al., 2021) can clearly resolve the CLGBs crossing the internal capsule connecting the caudate nucleus and putamen. In addition, the CLGBs are characterized by a high T2 weighted signal and a low fractional anisotropy (FA) in the range of values associated with other gray matter (GM) structures such as the cortex (FA ∼0.15 (Little and Beaulieu, 2021)). The sensitivity of MRI and diffusion MRI (dMRI) to CLGBs suggests that these imaging modalities are useful for studying these structures in-vivo in humans. Recent work has used in-vivo T1 weighted 3D fast spoiled gradient-echo images acquired at 0.9 mm isotropic to characterize the volumetric properties of CLGBs in a cohort of patients with normal or near-normal brain imaging findings, reporting an average CLGB thickness and length of ∼1.0 mm and ∼4.6 mm respectively (Dang et al., 2023). To our knowledge, this is the only previous study using neuroimaging to quantify properties of CLGBs in-vivo.

**Figure 2:**
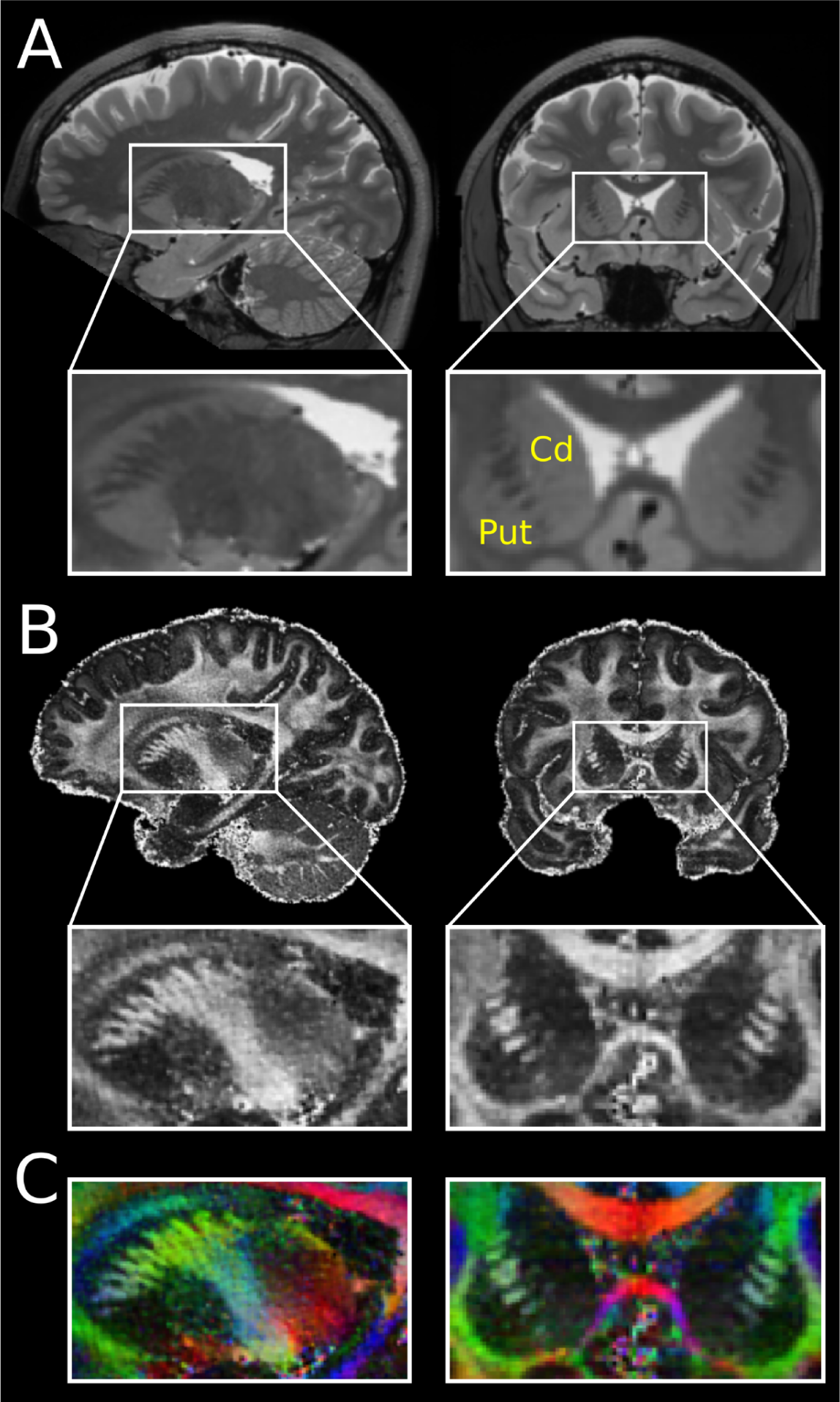
Visualization of the macroscopic caudolenticular gray matter bridge (CLGB) anatomy visible in high resolution magnetic resonance imaging. Anatomically, the CLGBs connect the putamen (Put) and the caudate nucleus (Cd) and are oriented orthogonally across the internal capsule. Example images depicting CLGBs are shown for a single subject acquired on (A) high-resolution T2-weighted imaging (0.75 mm isotropic) and (B) a fractional anisotropy map and (C) a fractional anisotropy map colored by primary diffusion direction generated from the diffusion imaging dataset acquired at 0.76 mm isotropic (Wang et al., 2021).

Diffusion MRI (dMRI) is an imaging technique sensitive to the movement of water molecules within brain tissue. White matter (WM) structural connectivity can be reconstructed between brain regions using streamlines tractography by traversing fiber orientation distribution function (fODF) maps generated from the dMRI signal (Jeurissen et al., 2019). Although dMRI and tractography are typically considered techniques for studying the microstructure and connectivity of WM, previous work has demonstrated that anisotropy of the diffusion signal can also be detected in the cortical GM (Calamante et al., 2018; McNab et al., 2013; Truong et al., 2014). More recent studies proposing the use of diffusion MR imaging acquired at a much higher resolution than conventional dMRI have reported reduced fractional anisotropy in regions associated with the CLGBs in both ex-vivo marmoset (Saleem et al., 2023) and in-vivo human (Wang et al., 2021) experiments but the orientation of diffusion anisotropy in these regions was not discussed. In fact, using standard preprocessing and modeling parameters for fODF maps, anisotropy of the diffusion signal can be observed in regions corresponding to CLGBs (Figure 3). In dMRI data acquired as part of the HCP (multishell b1000, b2000, b3000, 1.25 mm isotropic) preprocessed using standard methods (Theaud et al., 2020), fODF peaks oriented orthogonally across the internal capsule can be observed between the caudate nucleus and the putamen (Figure 3A). These “secondary” fODF peaks form a crossing with the larger amplitude “primary” fODF peaks belonging to the WM of the internal capsule. When plotted, the relative amplitude of the secondary peaks clearly delineate a striate pattern with larger peak amplitudes observed in distinct regions but not the entirety of the striatal zone, suggesting that these peaks correspond to the “bridges” of the CLGBs (Figure 3B). Given the small size of the fODF lobes corresponding to the CLGBs, it is unclear whether these peaks continuously follow the CLGBs along the full medial-lateral extent of these structures. By reconstructing streamlines along the CLGBs, streamlines tractography is a potential method to confirm that the “small” fODF peaks follow the CLGBs and the resulting streamline reconstructions would provide the local orientations necessary to extract a microstructural measurement from the CLGBs themselves. However, standard local probabilistic tractography algorithms designed for whole brain WM tractography only reconstruct a small number of CLGB-streamlines traversing these secondary peaks because 1) a small portion of tractography seeds are initiated within the internal capsule and 2) the probabilistic sampling of fODFs during streamline reconstruction will heavily favor the larger fODF peaks corresponding to the internal capsule WM (Figure 3C). Importantly, even though the terminology of diffusion MRI tractography is often associated with WM fibers (e.g. streamlines, “fiber” ODF), the streamlines reconstructed throughout this work traverse the fODF peaks associated with the GM of the CLGBs which are microstructurally different than primarily myelinated WM fibers. Thus, the terms CLGB-streamline, streamline and fiber used throughout should not be interpreted to describe the underlying CLGB microstructure, but rather the terms are retained for consistency with their previously described methodological meanings.

**Figure 3.**
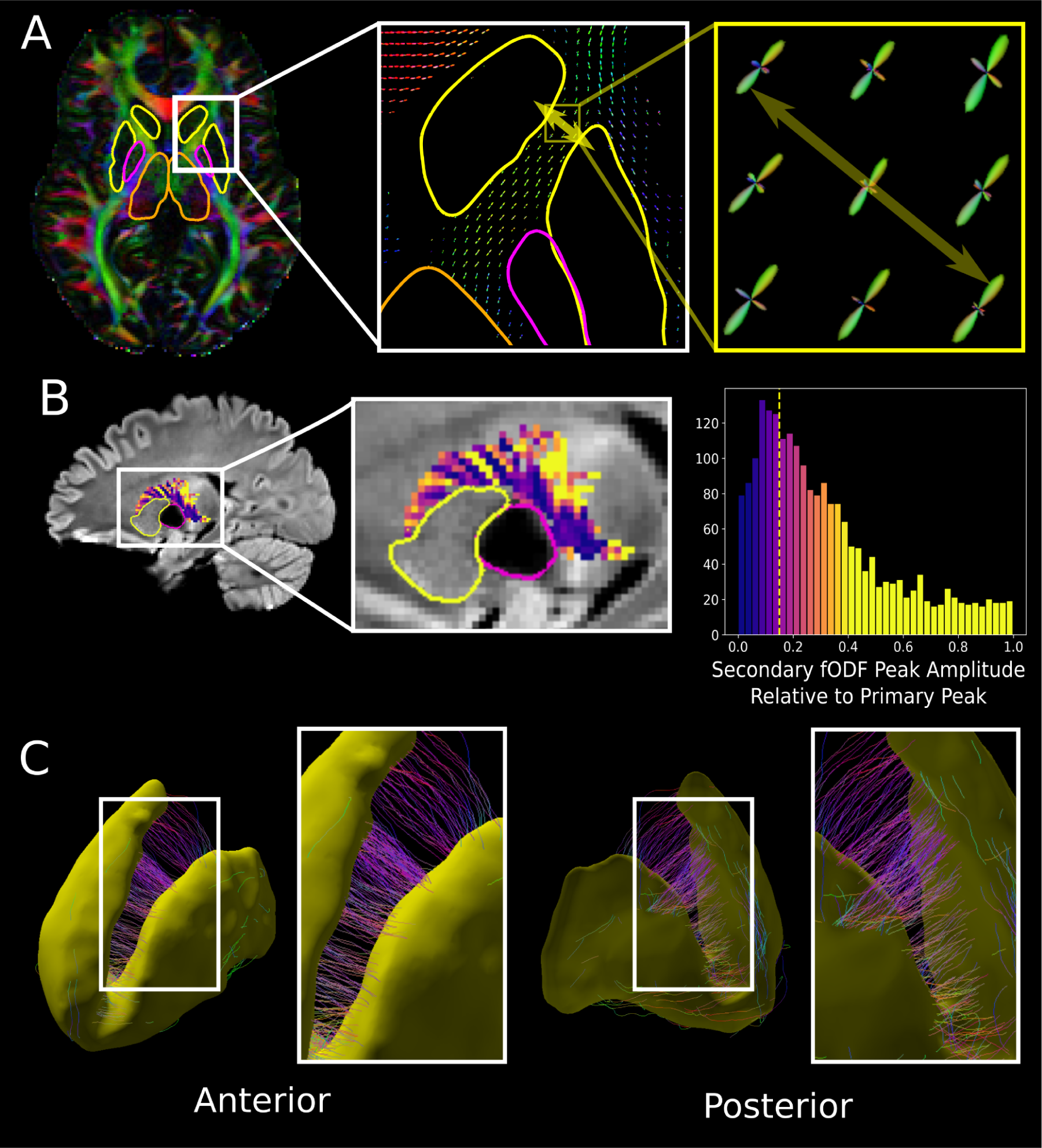
Secondary fODF peaks and standard tractography reconstruction across the internal capsule generated with a standard method and data acquired for a single subject from the HCP test-retest cohort acquired at 1.25 mm isotropic. (A) An axial RGB FA map shown with surface intersections of the striatum (yellow), globus pallidus (pink) and the thalamus (orange) visualized alongside fODF maps (zoomed images) depicting “secondary” fODF peaks oriented orthogonally across the internal capsule. (B) The relative peak amplitude values (secondary peak amplitude / max peak amplitude) extracted in a manually drawn internal capsule ROI are overlaid on a mean diffusion weighted b1000 image alongside a histogram of the values. A striate pattern in the internal capsule is visible in the relative peak values. (C) Results from a standard whole brain tractography pipeline that randomly seeded 1 million streamlines within white matter and did not use a streamline length threshold. After filtering to include streamlines with both ends in the striatum and less than 20 mm in length, few CLGB-streamlines are retained.

The small amplitude of these secondary fODF peaks and the difficulty of tracking these specific connections likely explains the lack of reference to CLGB-streamlines in the tractography and/or diffusion MRI literature. To date, few tractography studies have attempted to track between subcortical gray matter structures, and are focused on reconstructing WM pathways. In order to reconstruct WM pathways connecting the basal ganglia, studies have used standard whole brain tractography approaches (Kai et al., 2022; Worbe et al., 2015), targeted tractography methods designed for specific WM pathways implicated for neurosurgical applications (Avecillas-Chasin and Honey, 2020; Rozanski et al., 2017) or advanced super-resolution imaging techniques (Kwon et al., 2021), with no studies reporting the reconstruction of CLGB-streamlines. Given the absence of CLGB-streamlines in previous studies that have used standard tractography approaches as well as the demonstrated utility of tract specific approaches to adequately reconstruct short-range WM tracts crossing the internal capsule, it can be inferred that a more targeted tractography algorithm is needed to efficiently reconstruct CLGBs.

Here, a novel tractography seeding approach is proposed that incorporates the geometry of the striatum based on a 3D surface reconstruction of the structure. Previous work has proposed using the surfaces of subcortical GM structures for filtering (Marrakchi-Kacem et al., 2013) and creating anatomical stopping criteria (Yeh et al., 2017) for reconstructing cortical to subcortical fiber projections. Other work has proposed using the WM/cortex interface surface to overcome the gyral tractography bias (St-Onge et al., 2018). To reconstruct CLGBs, a novel surface-based tractography seeding strategy is proposed that seeds streamlines orthogonally outwards from the GM/WM interface of the striatum. The method is used to reconstruct CLGB-streamlines that overlap expected high T2-weighting and low FA GM regions on 760 µm isotropic diffusion data. By downsampling this data, the novel seeding method is shown to be capable of reconstructing CLGBs at more readily acquired diffusion imaging resolutions. Repeatability of the CLGB reconstruction method is assessed on two test-retest cohorts; the HCP test-retest dataset acquired at 1.25 mm isotropic and a standard diffusion imaging test-retest cohort acquired at 2 mm isotropic. Finally, a first attempt at measuring diffusion anisotropy along the CLGBs is presented by leveraging the CLGB-streamline orientations and density output from the CLGB tractography reconstructions to measure the amplitude of the “secondary” fODF lobes associated with these GM structures.

## 2 Materials and Methods

### 2.1 Participants and Data Acquisition

The proposed method was evaluated on two publicly available diffusion data sets and one previously published test-retest diffusion dataset, namely; a high resolution isotropic diffusion data acquisition (Wang et al., 2021) is used to demonstrate the spatial accuracy of the proposed CLGB reconstruction method and assess the effects of downsampling the image resolution, the test-retest cohort acquired as part of the human connectome project (HCP) (Glasser et al., 2016) and another test-retest dataset using a more standard diffusion acquisition was used to assess repeatability of CLGB reconstructions (Edde et al., 2023). The high resolution in-vivo data was acquired at 3T on a single subject at 760 µm isotropic resolution with multi-shell q-space sampling separated over multiple sessions. Only the first three sessions of the diffusion acquisitions were used, which each contained 14 b0 and 140 b1000 images after correcting for distortions with reverse phase encoding. This acquisition also acquired multiple T2 weighted images with the T2-SPACE sequence at 0.7mm isotropic. In the current study, the average of these three images is used as a reference for the location of CLGBs (further acquisition details can be found (Wang et al., 2021)).

The test-retest HCP cohort includes multimodal imaging data acquired at two different time points for 46 individuals and only those who had diffusion data available at both time points (N=44) were included in this study. The diffusion images for the HCP data were acquired at 1.25mm isotropic resolution with all shells (b0, b1000, b2000 and b3000) used in this study. Data was preprocessed using the standard HCP diffusion processing pipeline outputting eddy current corrected diffusion weighted images, gradient values and whole brain masks used for subsequent analysis. T2-weighted structural imaging acquired at 0.7 mm isotropic was also used in the current study as a reference for CLGB locations in each subject/session (further acquisition details can be found (Glasser et al., 2016)).

The “clinical” test-retest dataset was acquired serially (up to 5 repeat scans on separate visits) on multiple subjects (further details can be found (Edde et al., 2023)). In this study, only images from 24 subjects that had completed T1-weighted imaging and diffusion imaging for the two first imaging sessions were used. Whole brain MRI images were acquired on a clinical 3T MRI scanner (Ingenia, Philips Healthcare) using a 32-channel head coil. Whole brain T1 images were acquired axially using a 3D T1-weighted MPRAGE sequence at 1.0 mm isotropic resolution, TR =7.9 ms, TE = 3.5 ms, TI = 950 ms, FOV = 224 x 224 mm^2^ yielding 150 slices, flip angle = 8 degrees, with a total acquisition time of 4 min 20s. Multishell whole brain diffusion images were acquired with a single-shot EPI spin-echo sequence at 2.0 mm isotropic resolution, TR = 4800 ms, TE = 92 ms, SENSE factor = 1.9, Multiband-SENSE factor = 2, flip angle = 90 degrees, FOV = 224 x 224 mm^2^, over 66 slices, with a total acquisition time of 9 min 19s. Diffusion weighting included multiple shells with 100 unique diffusion directions uniformly distributed over each shell (b300 8 directions, b1000 32 directions, b2000 60 directions), 7 b0 images and a reverse phase encoded b0 image. For this study to simulate a practical “clinical” acquisition only the b0 and b1000 images were used and the reverse phase encoded image was omitted (not typically acquired in clinic), which corresponds to a total acquisition time of ∼3.5 min compared to the total acquisition time of the entire multishell acquisition of 9 min 19 seconds.

### 2.2 Surface Based Brain Segmentation and White Matter Mask

Segmentations of subcortical and cortical brain structures were generated using 3D deformation methods (Little et al., 2023; Little and Beaulieu, 2021) using only the b0 and b1000 diffusion weighted images. This segmentation approach uses the image contrast visible on the powered average b1000 diffusion image, FA and mean diffusivity (MD) maps to identify the boundaries of subcortical and cortical structures, outputting 3D triangular meshes (surfaces) consisting of vertices and edges for the left/right globus pallidus, striatum, thalamus, hippocampus, amygdala, WM/cortex boundary and cortex/cerebrospinal fluid (CSF) boundary. This segmentation approach was chosen because 1) the boundaries of the outputted 3D surface lie at the WM/GM interface visible on the diffusion images (i.e. no potential registration errors to T1w imaging), thus tractography streamlines can be seeded directly from the surface vertices located at the interface, and 2) the 3D surface can be used to compute angles to inform the initial seed direction for streamlines. In addition to GM surfaces, an accurate WM mask used for streamlines tractography was generated by including all voxels within the WM/cortex boundaries and excluding all voxels within the subcortical GM and lateral ventricles.

### 2.3 Caudolenticular gray matter bridge reconstruction using subcortical surface based tractography seeding and filtering

To reconstruct the CLGBs and to avoid tracking WM fiber bundles that dominate the diffusion signal (i.e. primary fODF peaks) within the internal capsule, a tractography seeding and filtering strategy is proposed that utilizes the 3D geometry of the striatum. A workflow for the proposed method is visualized in Figure 10. First, rather than seeding from random locations within the white matter mask, streamlines are initiated from the location of each vertex of the 3D striatum surface (Figure 10A). To encourage an even distribution of seeds across the surface, prior to seeding, the surface is resampled such that the distance between vertices is approximately 0.1 mm. To identify vertices bordering the internal capsule an ROI was generated by first registering the JHU white matter atlas to the subject’s native imaging space with ANTS SyN registration (Avants et al., 2008). The transformation was calculated by registering the HCP FA template (FSL v6.0) to the subject’s FA map and the JHU atlas was resampled to the subject’s native imaging space using a nearest neighbors interpolation. Then, a left and right seed selection ROI is generated by including all WM labels adjacent to the caudate nucleus and putamen. This ROI was then dilated using a 2 mm sphere and vertices included in the dilated ROI are identified as bordering the internal capsule (Figure 4A). Streamlines are then initiated from each identified vertex in the direction outward and normal to the striatum surface. Standard probabilistic tractography is performed from each seed 10 times using a spherical function threshold of 0.15 (0.10 for the clinical dataset) keeping all streamlines (regardless of length), while using all other default tracking parameters (Theaud et al., 2020) generating a striatum specific tractogram representing projections outward from the 3D surface of the striatum (Figure 4B).

**Figure 4.**
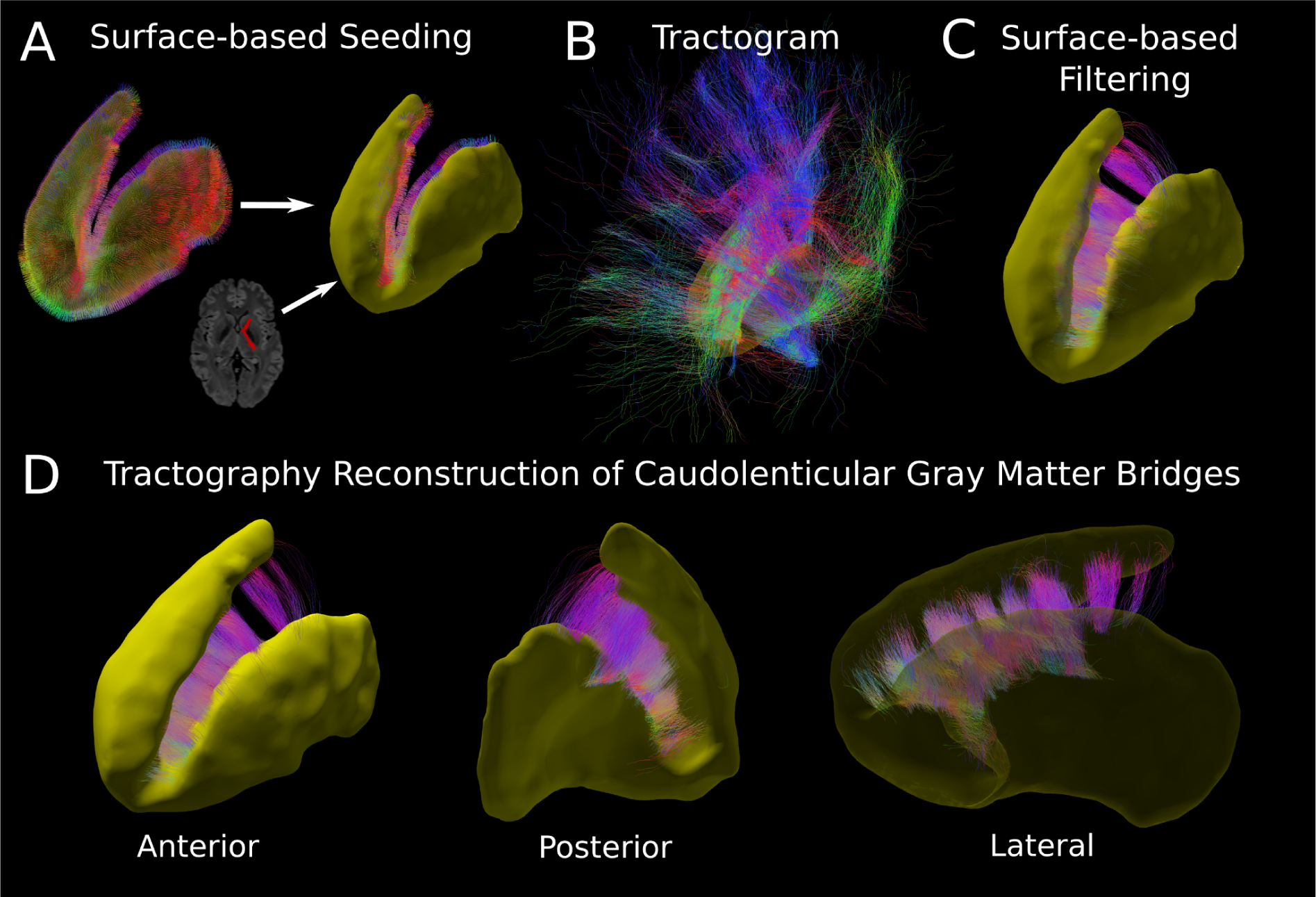
Workflow for the proposed tractography method for reconstructing CLGBs. (A) Based on the 3D surface of the striatum (yellow) surface normals are generated at each vertex location. Then vertices adjacent to the internal capsule are identified based on a registered atlas ROI (red) and these vertices/normals are retained for tractography seeding. (B) Standard local probabilistic tractography is performed starting at the identified vertices setting the initial search cone orthogonal to the striatum surface. (C) The tractogram is filtered including streamlines with both ends in the striatum that are less than 20 mm in length. (D) The resulting CLGB reconstruction is visualized in 3D for the left striatum for an example subject depicting a dense reconstruction of CLGB-streamlines crossing the internal capsule.

To filter tractograms such that only CLGB-streamlines remain, surface-based proximity filtering is proposed to remove streamlines based on endpoint proximity to the boundary of the target 3D surface. In other words, if the distance from the surface to a streamline’s endpoint is inside or within a distance threshold of the target surface then the streamline is retained. In this case, the striatum specific tractogram is filtered first to include streamlines with both ends inside the striatum surface (distance threshold 0.1 mm) generating a CLGB reconstruction that only contains CLGB-streamlines between the caudate and the putamen. To retain only streamlines that are associated with the CLGBs, resulting tractograms are then filtered to exclude a small proportion of streamlines that are either greater than 20 mm in length or travel greater than 6 mm in the posterior-anterior direction(Figure 4C and 4D).

### 2.4 Caudolenticular gray matter bridge reconstruction at submillimeter resolution

To demonstrate the ability of the proposed method to reconstruct CLGBs, the method was first applied to the high-resolution diffusion data acquired at 760 µm isotropic. Firstly, the raw diffusion data for each imaging session containing b0 and b1000 diffusion weightings (i.e. session 1, session 2, session 3) was denoised (Veraart et al., 2016) (DIPY version 1.7.0). For each session, subcortical structures and the inner WM/GM cortex boundary were segmented and 3D surfaces along with a white matter tracking mask were output as previously described. Then CLGBs were reconstructed for each imaging session using the proposed tractography seeding and filtering method. To ensure reconstructions were accurately capturing CLGBs visible in other imaging maps, CLGB-streamline intersections were overlaid on the average T2-weighted structural image (registered to each session, T2 image to mean b0 image, rigid body transformation (ANTs Version 2.3.5)) and FA maps calculated for each session independently.

### 2.5 Evaluation of caudolenticular gray matter bridge reconstruction at multiple resolutions

To assess the effect of image resolution on CLGB reconstruction, the analysis was repeated at multiple downsampled resolutions using the 760 µm diffusion data as follows. Firstly, the denoised diffusion data for each session was resampled to coarser image resolutions (cubic spline interpolation) namely, 1.0 mm, 1.25 mm, 1.5 mm, 1.75 mm and 2.0 mm. Following downsampling, a white matter mask and fODF map were recalculated at each resolution as previously described. Then CLGBs were reconstructed using the proposed tractography seeding and filtering method, keeping all tracking parameters the same for each resolution. To isolate the effect of resolution, the same tractography seeds from the striatum segmentation generated at the original 760 µm resolution are used for all coarser resolutions from the same session. To investigate the relationship between T2 image intensity and CLGB reconstructions, streamline intersections were visualized on top of the T2 image for CLGBs reconstructed at each resolution.

To quantify the effect of resolution on this single subject, overlap metrics were calculated between the CLGB reconstructions generated at 760 µm and reconstructions at each downsampled resolution for each session. For each downsampled resolution Dice score, weighted Dice score (Cousineau et al., 2017), volume overlap and volume overreach were calculated. Here, Dice score is calculated as two times the number of voxels intersected by both reconstructions divided by the sum of total voxels intersected by reconstruction one and total voxels intersected by reconstruction two. Weighted Dice score is an extension of Dice score that incorporates the streamline density at each voxel to weight more densely traversed regions in the Dice score measure (Cousineau et al., 2017). Both weighted Dice score and Dice score range from 0 to 1, where 1 indicates a perfect overlap between streamline reconstructions and 0 indicates no overlap between reconstructions. Volume overlap is the total volume intersected by both streamline reconstructions divided by the CLBG volume at 760 µm and volume overreach is the total volume intersected by a single streamline reconstruction divided by the CLGB volume at 760 µm. Each metric was averaged across each session for each downsampled resolution keeping the left and right hemispheres separate.

### 2.6 Repeatability of Caudolenticular Gray Matter Bridge Reconstruction in Test-Retest Cohorts

The proposed seeding approach was performed on each subject and session from the HCP test-retest cohort and “clinical” 2mm isotropic test-retest cohort generating left/right CLGB reconstructions for each subject and session. To assess the repeatability of the subcortical GM connections reconstructed by the proposed tractography seeding strategy, a rigid transformation from session 2 to session 1 was calculated using the mean b0 image from the scans (ANTs Version 2.3.5). The transformation was then applied to the CLGB-streamlines for each hemisphere. To quantify repeatability, Weighted Dice score, Dice score (Cousineau et al., 2017), Volume Overlap and Volume Overreach were calculated for each subject and across sessions (scan 1 used as reference for volume overlap and overreach). All metrics were averaged across hemispheres for each session.

### 2.7 Fixel-based measurements of “secondary” fODF peaks associated with CLGB

To measure the lobe characteristics associated with the CLGBs and to avoid the measurement of the larger fODF peaks associated with the WM of the internal capsule, a fixel-based measurement of lobe amplitude is combined with weighting from the streamline density maps. First for each reconstructed CLGB, normalized streamline density maps (# streamlines per voxel divided max density) were generated. Next, fODF lobe metrics are generated using a bingham fit (Riffert et al., 2014) followed by calculating the integral of the bingham function on the sphere for each fODF lobe (https://github.com/scilus/scilpy/blob/master/scripts/scil_fit_bingham_to_fodf.py). Fixel-based fODF lobe integral maps were then output by calculating the fODF lobe integral for each CLGB-streamline intersecting a voxel and then assigning the average lobe integral across streamlines to that voxel. The bingham fit fODF lobe integral was chosen because it is a more accurate measure of fODF lobe geometry compared to the more commonly referred to apparent fiber density (Riffert et al., 2014).

Then a single weighted average is calculated for each CLGB reconstruction for each hemisphere by averaging all non-zero voxels in the fODF lobe integral map weighted by the normalized streamline density map. Four comparison metrics were attained, namely; CLGB volume calculated for each reconstruction; mean FA (average of all voxels with at least one CLGB-streamline); streamline weighted mean FA (average FA weighted by the normalized streamline density map); mean fODF lobe integral (average of all voxels with at least one CLGB-streamline). All metrics were calculated for each subject in the 760 µm dataset (all 3 sessions), the HCP test-retest cohort (first scan only) and the “clinical” 2 mm dataset (first scan only). Metrics were averaged within each cohort keeping the left and right hemisphere separate and then hemispheric differences were tested with a paired t-test (p < 0.05) within each cohort per measure.

### 2.8 Quantitative comparison to other tractography seeding approaches

As a comparison to the proposed surface-based seeding strategy, tractograms were reconstructed using two other comparable tractography seeding methods on the HCP test-retest data (first scan only). Firstly, tractograms were created for the striatum by seeding streamlines at each surface vertex as in the proposed surface-based method but randomly sampling an initial direction from the fODF rather than explicitly setting the initial direction based on the surface normal. Secondly, a more standard tractography approach was used by seeding randomly within an internal capsule ROI. In this instance the internal capsule ROI consists of the posterior and anterior internal capsule labels extracted from the previously registered JHU atlas. As in the original experiment, tractograms from the two comparable approaches were filtered using surface-based distance filtering as well as length and orientation criteria. Reconstructions were generated for both comparable methods for each hemisphere for all 44 subjects from the HCP test-retest cohort using the first scan only. As a measure of how efficiently each method performed in reconstructing CLGB connections, the number of CLGB-streamlines generated by each method were calculated per hemisphere for each subject. As a measure of voxel coverage for each method, the average CLGB-streamline density and percentage of voxels with at least one CLGB-streamline were calculated within the internal capsule ROI per hemisphere for each subject. Group average and standard deviations were calculated for each output measure, and pairwise t-tests were conducted (p < 0.05) to assess differences between the proposed method and alternative approaches for each output measure.

## 3 Results

### 3.1 Validation of CLGB reconstruction method at a submillimeter resolution

The proposed surface-based seeding CLGB reconstruction method was successfully repeated on all 3 sessions of the 760 µm diffusion imaging dataset generating reconstructions in all cases. Visualization of the CLGB reconstructions for the 3 sessions are shown in Figure 5, along with streamline intersections overlaid on the T2-weighted image and FA maps from each session. Streamline intersections from left and right CLGB reconstructions clearly intersect regions associated with a high T2 weighted signal and low FA in the range of GM values, indicating that the CLGB-streamlines are traversing fODF peaks corresponding to GM and not WM. In addition, the spatial distribution of CLGB-streamlines is consistent across all sessions suggesting the same regions are being traversed by the proposed reconstruction method.

**Figure 5.**
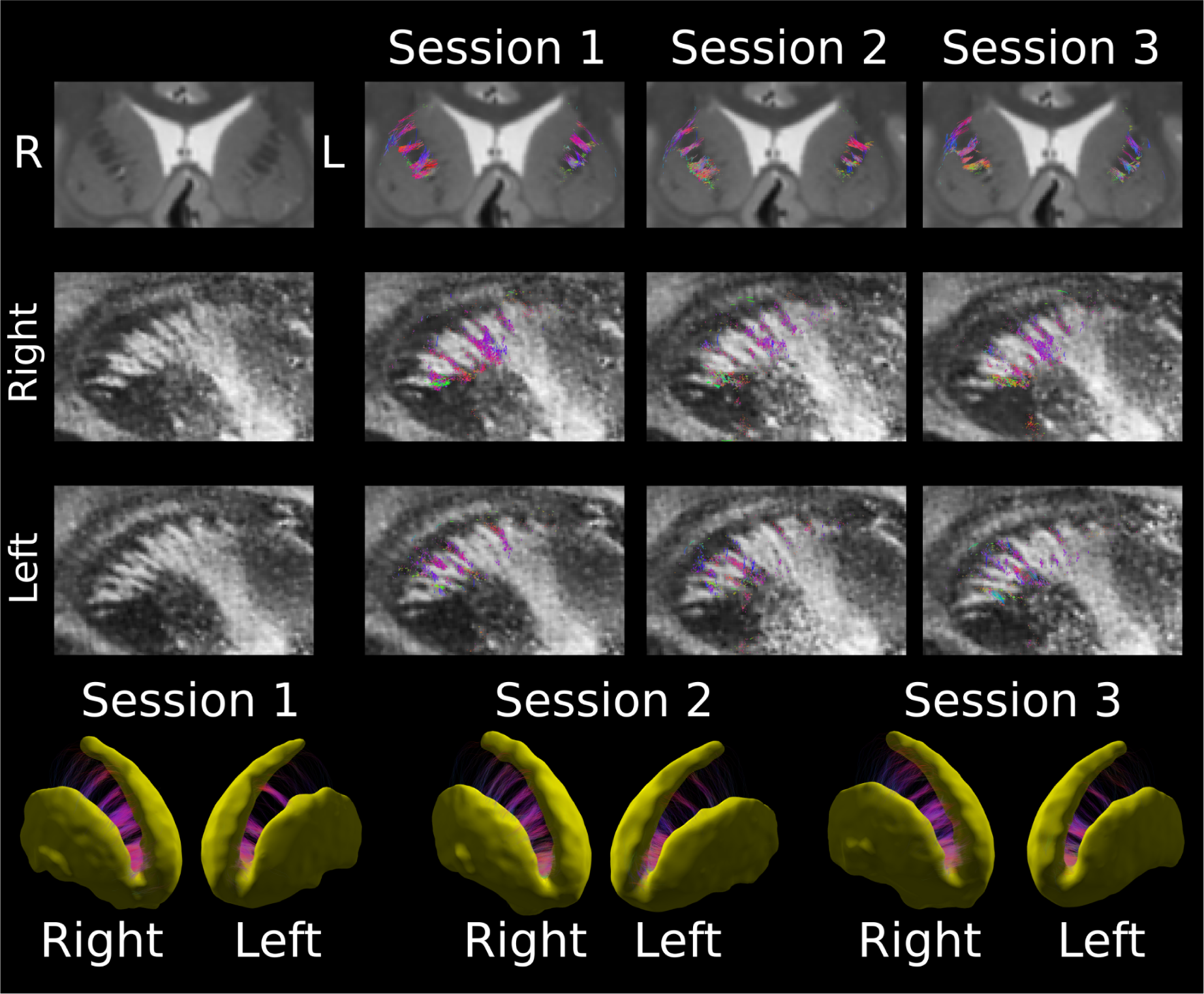
Caudolenticular gray matter bridge (CLGB) tractography reconstructions for the same subject generated from 3 sessions from the 760 µm isotropic diffusion dataset. CLGB reconstructions are visualized as streamline intersections on a coronal slice from a T2-weighted structural image registered to each session (row 1) and on a sagittal slice from the fractional anisotropy maps for the right (row 2) and left (row 3) hemispheres. CLGBs and striatum segmentations (yellow) generated for each session are displayed in row 4. Reconstructions for CLGBs consistently intersected regions of relatively high T2 signal intensity as well as low fractional anisotropy and were distributed similarly across all sessions.

### 3.2 Effect of image resolution on CLGB reconstruction

Using the proposed method and the 760 µm isotropic diffusion dataset, CLGB tractograms were reconstructed on both hemispheres on interpolated coarser resolution diffusion data (i.e. 1.0, 1.25, 1.5, 1.75 and 2.0 mm isotropic). The CLGB reconstructions are visualized in Figure 6 as 3D renderings and as slice intersections on the T2 weighted structural image acquired at 0.75 mm isotropic (interpolated to 0.5 mm isotropic for better visualization of CLGBs). Notably, the reconstructions at 760 µm largely overlap the expected locations of CLGBs visible on the T2 weighted image. As the resolution becomes more coarse, the correspondence between the T2 image intensity and the CLGB reconstructions diminishes. Additionally, at the coarsest resolutions (1.75 and 2.0 mm) the method fails to reconstruct the thin CLGBs connecting the posterior portion of the striatum surfaces but the anterior CLGB-streamlines are retained. Whereas at finer resolutions (1.0, 1.25 and 1.5 mm) thinner CLGBs are reconstructed but appear visibly thicker due to signal blurring.

**Figure 6.**
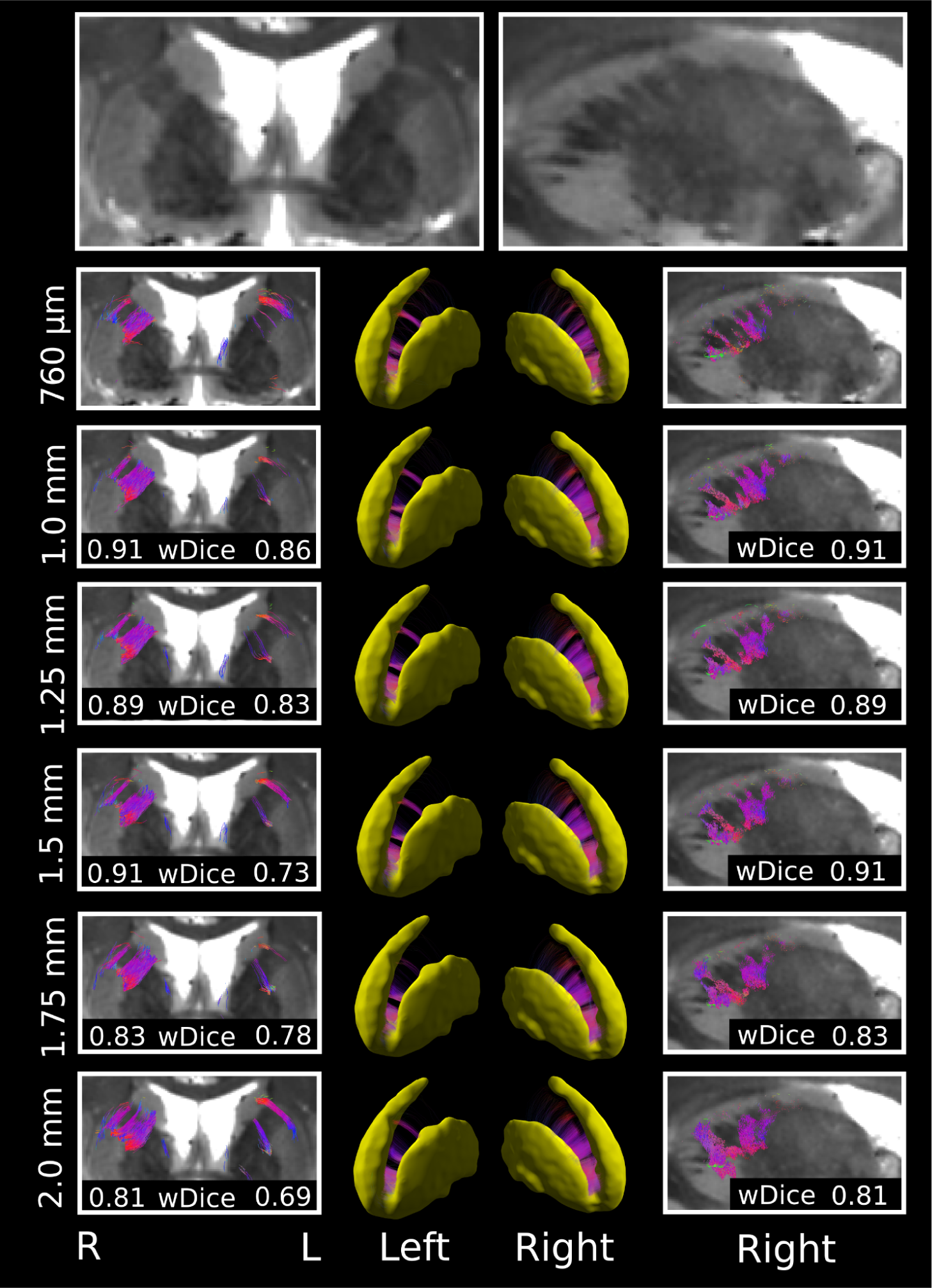
Tractography reconstruction of the left/right CLGBs generated using the high resolution (760 µm isotropic) diffusion dataset. The proposed tractography method was performed at the native resolution and then repeated at multiple coarser resolutions (1.0, 1.25, 1.5, 1.75, 2.0 mm isotropic) by interpolating the diffusion data set. Tractograms are viewed as 3D renderings and streamline intersections overlaid on a T2 weighted image acquired at 0.75 mm isotropic (interpolated to 0.5 mm isotropic to better visualize the CLGBs). Streamline weighted Dice scores (wDice) calculated between the downsampled CLGB reconstructions and the reconstruction generated at the native 760 µm are also displayed for the left and right hemispheres at each downsampled resolution. Reconstructions of the anisotropic CLGBs were generated at all downsampled resolutions. At more coarse resolutions the overlap (wDice) with the native resolution reconstruction gradually degraded, with thinner bridges failing to be reconstructed at the coarsest resolutions (1.75 and 2.0 mm).

Quantitatively, effects of resolution were assessed using overlap metrics calculated between the interpolated coarser resolution CLGB reconstructions and the CLGB reconstructions generated at the native imaging resolution. Average metrics across the three sessions are shown in Figure 7. Weighted Dice score and Dice score decreased as a function of resolution, whereas volume overlap stayed relatively constant across resolutions (left: ∼72% at 1.0 mm to ∼58% at 2.0 mm, right: ∼76% at 1.0 mm to ∼ 62% at 2.0 mm) compared to volume overreach which increased at lower resolutions (left: ∼111% at 1.0 mm to ∼144% at 2.0 mm, right ∼70% at 1.0 mm to 93% at 2.0 mm). Taken together these results suggest that at lower resolutions CLGB reconstructions overlap regions of CLGBs visible at higher resolutions but are distributed across more voxels due to spatial blurring of the signal.

**Figure 7.**
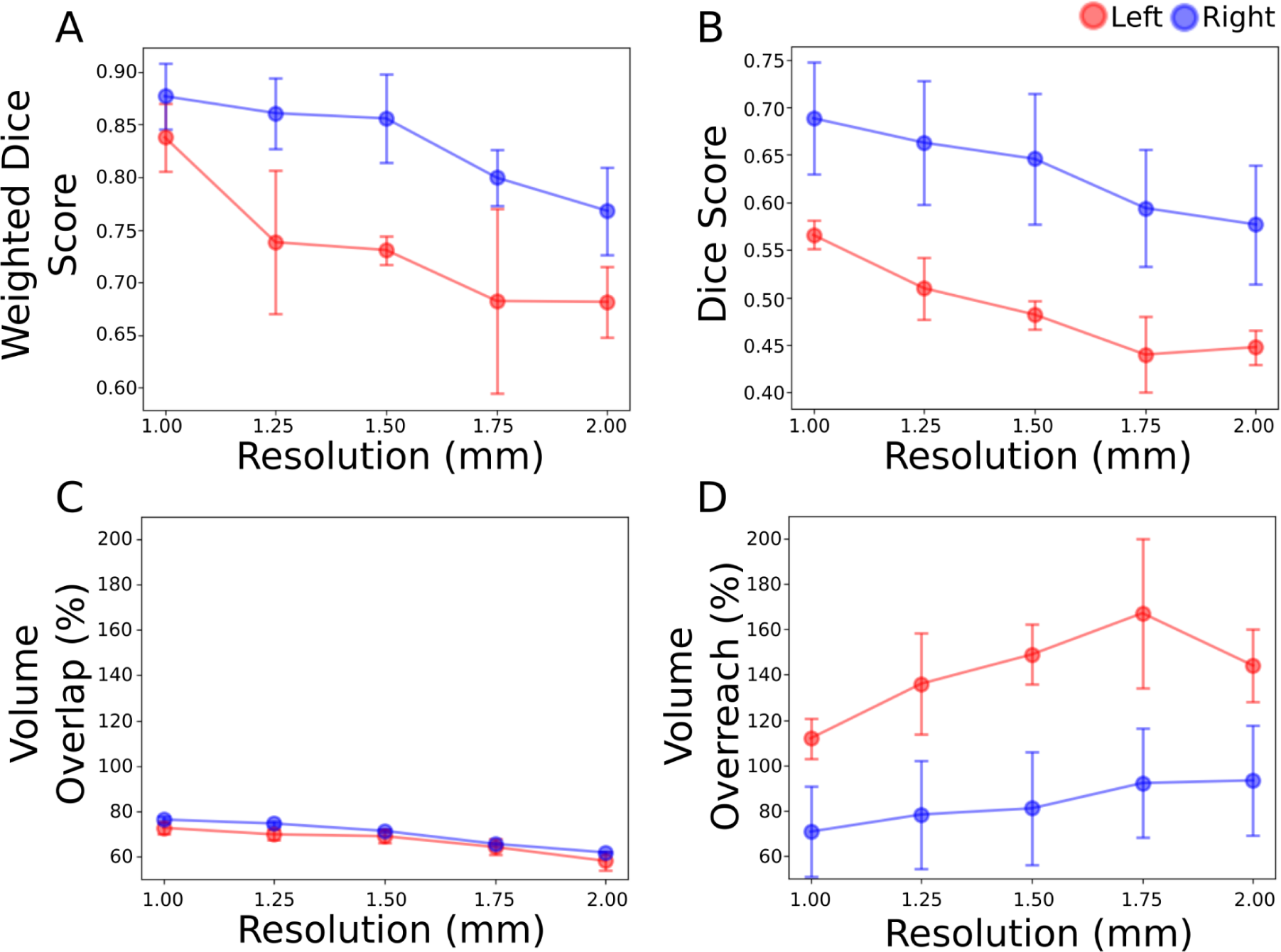
Average overlap measures between 760 µm CLGB reconstruction and CLGB reconstructions generated at different downsampled diffusion image resolutions. Weighted Dice score (A) and Dice score (B) both show lower values as a function of image resolutions, whereas volume overlap (C) had relatively stable values across image resolutions compared to volume overreach (D) which had higher values as a function of image resolution.

### 3.3 Test-retest repeatability of CLGB reconstruction

The proposed method was performed on all subjects and scans from the HCP test-retest cohort and the “clinical” test-retest cohort. CLGBs were reconstructed successfully in all subjects/scans from the two datasets. An interscan comparison of CLGB reconstruction is visualized in Figure 8 for a single subject from each cohort. Even in the presence of striatum surface-based segmentation differences, CLGBs reconstructed on session 1 and session 2 show marked similarity in both 3D renderings and streamline intersections overlaid on the T2 weighted structural images for the HCP data and a T1-weighted image on the clinical data. Quantitative test-retest metrics are reported in Table 1. Interscan weighted Dice scores calculated with session 1 and session 2 streamline density maps were noticeably high for the left and right CLGB reconstructions on both the HCP and clinical cohorts (weighted Dice > 0.93), meaning that regions with a high CLGB-streamline density were shared between session 1 and session 2. Dice scores for CLGBs had lower values (ranging from 0.72 to 0.80 between cohorts) relative to the weighted version of the same metric, suggesting that spurious fibers were the source of the discrepancy between measures.

**Figure 8.**
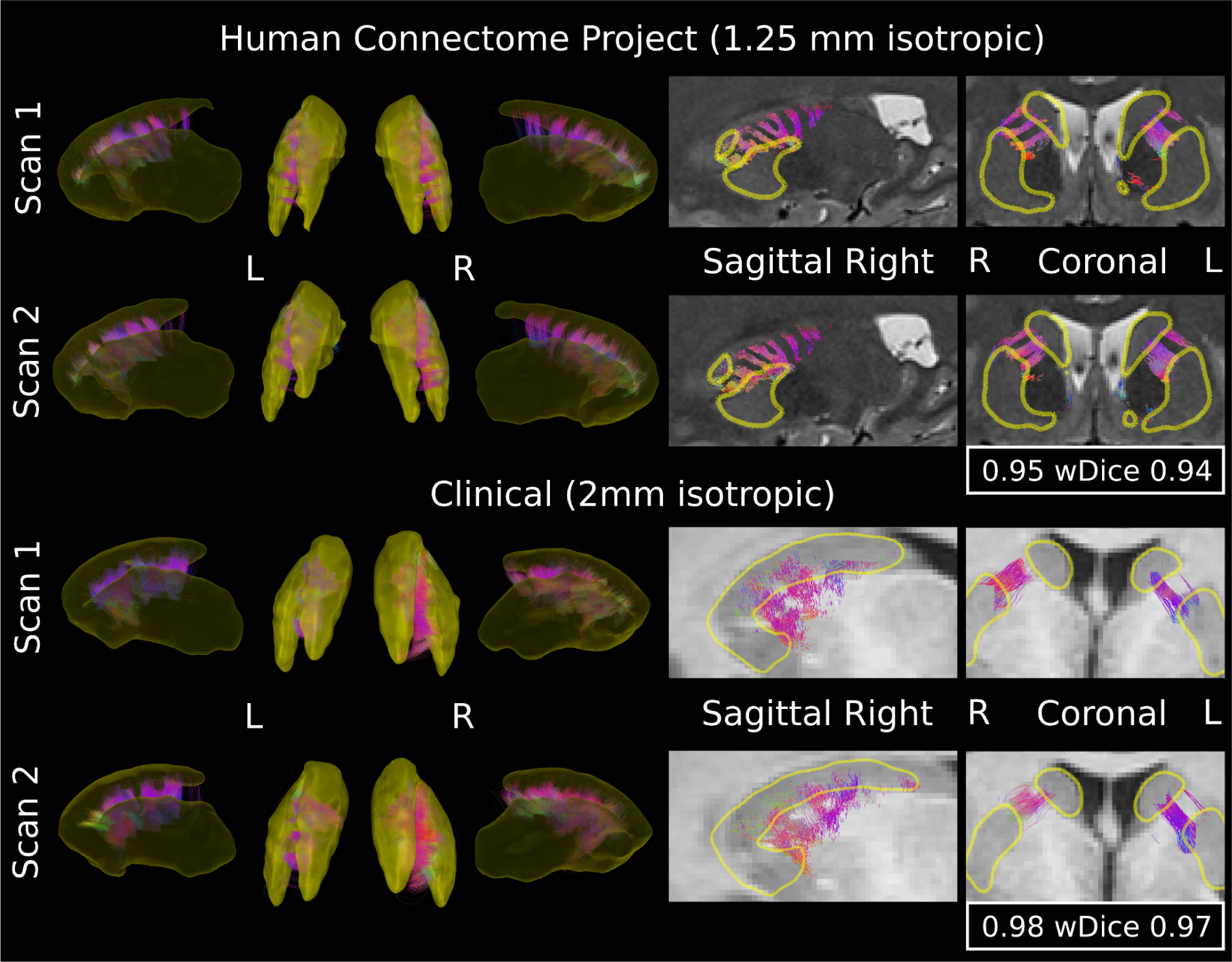
Visualization of caudolenticular gray matter bridge reconstructions on a single subject from the HCP test-retest cohort (scan 1: row 1, scan 2: row2) and a separate subject from the clinical test-retest cohort (scan 1: row 3, scan 2: row 4). Reconstructions are displayed as 3D renderings of the CLGB-streamlines between left and right striatum surfaces (yellow). Streamline intersections taken from approximately the same slice of the structural images (T2 weighted for HCP dataset, T1 weighted for clinical dataset) are displayed (column 4 and 5). Even with slight differences in test-retest segmentation, marked interscan similarity is observed in both visualizations corresponding to high interscan weighted Dice scores (wDice) for both the HCP (left: 0.94, right: 0.95) and clinical (left: 0.97, right: 0.98) datasets in these two examples.

**Table 1.**
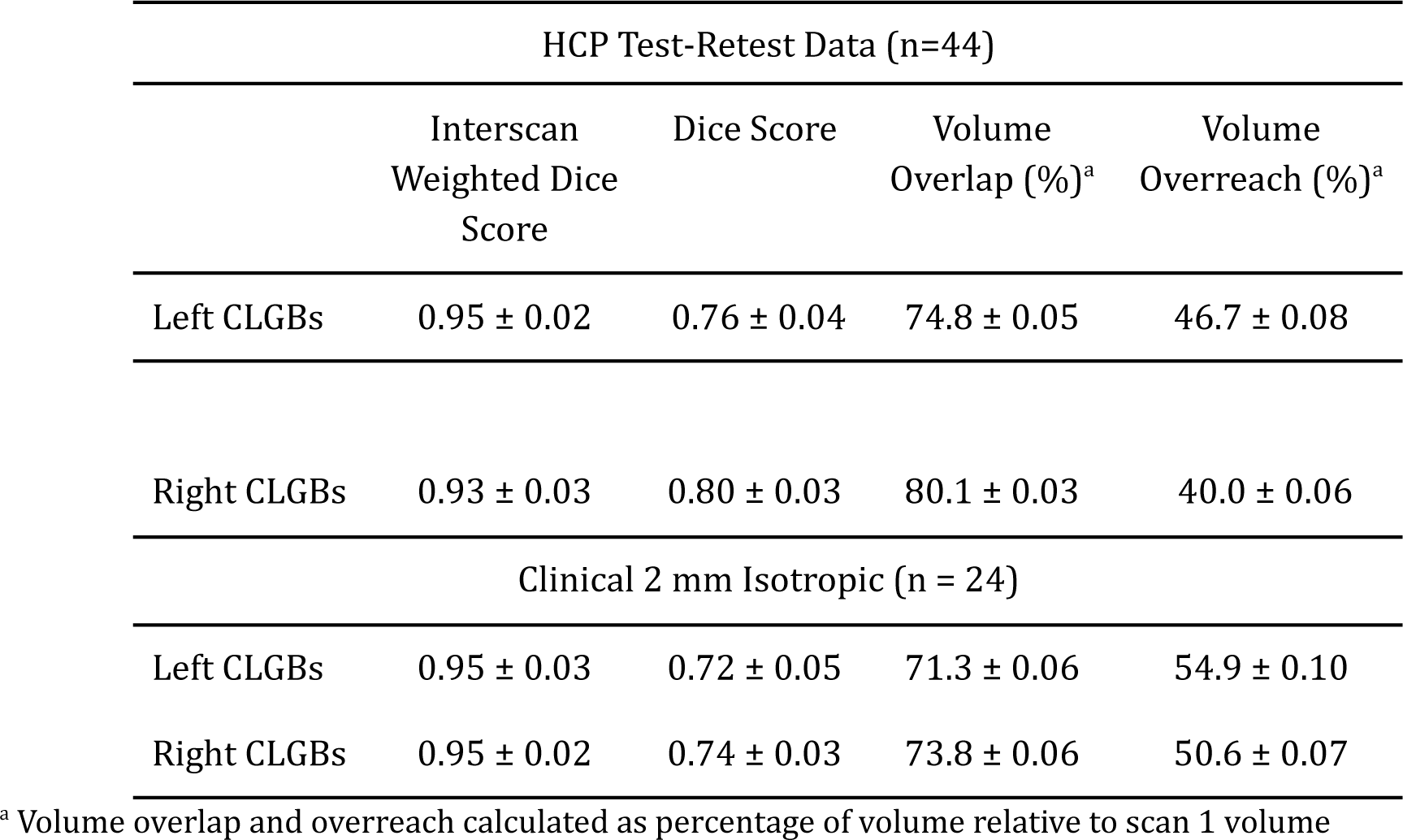
Measures of CLGB repeatability from the HCP test-retest and “Clinical” cohorts.

| HCP Test-Retest Data (n=44) |  |  |  |  |
| --- | --- | --- | --- | --- |
|  | Interscan<br>Weighted Dice<br>Score | Dice Score | Volume<br>Overlap (%)<sup>a</sup> | Volume<br>Overreach (%)<sup>a</sup> |
| Left CLGBs | 0.95 ± 0.02 | 0.76 ± 0.04 | 74.8 ± 0.05 | 46.7 ± 0.08 |
| Right CLGBs | 0.93 ± 0.03 | 0.80 ± 0.03 | 80.1 ± 0.03 | 40.0 ± 0.06 |
| Clinical 2 mm Isotropic (n = 24) |  |  |  |  |
| Left CLGBs | 0.95 ± 0.03 | 0.72 ± 0.05 | 71.3 ± 0.06 | 54.9 ± 0.10 |
| Right CLGBs | 0.95 ± 0.02 | 0.74 ± 0.03 | 73.8 ± 0.06 | 50.6 ± 0.07 |

<sup>a</sup> Volume overlap and overreach calculated as percentage of volume relative to scan 1 volume

### 3.4 Quantitative measurement of volumetric and diffusion properties of CLGBs

Maps of the fODF lobe integral along with normalized streamline density maps were calculated for the left/right hemisphere CLGB reconstructions for all subjects in the three cohorts used in the current study. Example quantitative maps are displayed in Figure 9 along with 3D renderings and slice intersections on a T2 image for CLGB reconstructions generated from a single subject from the HCP test-retest cohort. Both streamline density and the fixel-based lobe integral maps clearly show the same striate pattern visible in the CLGB-streamline intersection visualization. Higher lobe integral values were observed in regions of low CLGB-streamline density which likely reflects spurious streamlines on the edge of the CLGB reconstructions that intersect larger WM fiber bundles adjacent to the striatum. This supports the use of streamline weighting when extracting an average lobe integral value for the CLGBs.

**Figure 9.**
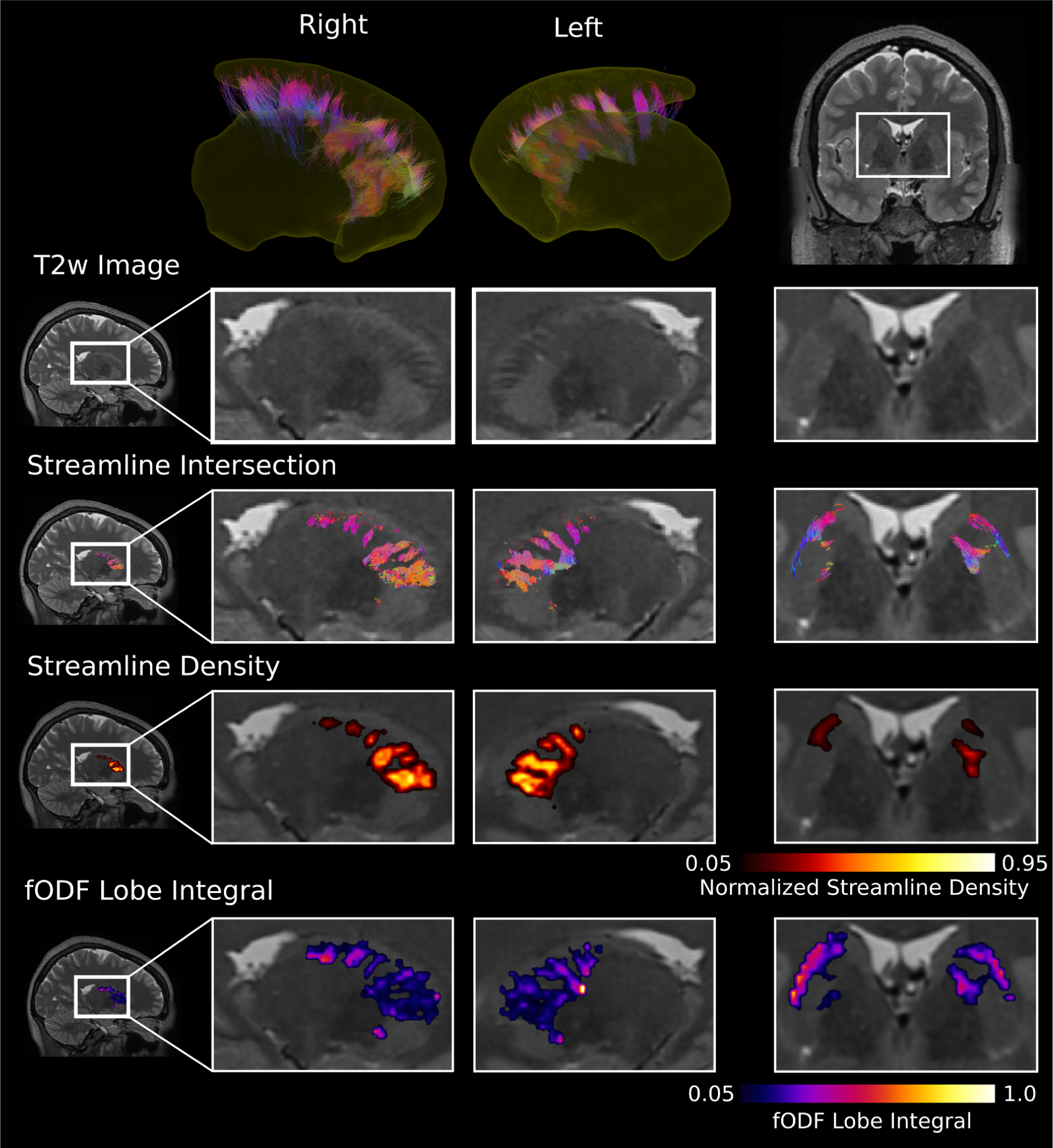
Tractography and quantitative maps for a left and right CLGB reconstruction for a single subject from the 1.25 mm isotropic HCP diffusion data. Maps and streamline intersections are overlaid on a 0.75mm isotropic T2 weighted structural image (interpolated to 0.5 mm to better delineate the CLGBs). Streamline intersection, streamline density and fODF lobe integral maps all overlap the CLGBs visible in the T2 weighted image. Lobe integral maps showed increased values in regions with low streamline density indicating these maps were sensitive to outlier streamlines bordering the CLGB reconstruction (column 3, row 3 and 4).

Quantitative metrics were extracted from the reconstructed left and right CLGBs, then averaged across each of the 3 cohorts and hemispheric differences were assessed. Average values for each cohort and t-test results are reported in Table 2. Higher volume was observed for right CLGBs compared to left CLGBs in all three cohorts. Unweighted diffusion metrics also demonstrated hemispheric differences in FA and lobe integral in all three cohorts. FA values were higher for the right hemisphere compared to the left in the high resolution and the clinical cohorts, but lower in the right hemisphere compared to the left hemisphere in the HCP cohort. A similar inconsistency between cohorts was observed with measurements of the fODF lobe integral which were higher for the left hemisphere compared to the right hemisphere in the high resolution data but lower in the left hemisphere compared to the right in the HCP and Clinical cohorts. Streamline weighted FA was higher for the left hemisphere in the HCP and Clinical data, whereas only a small hemispheric difference (left > right) in streamline weighted lobe integral was observed in the HCP data.

**Table 2.** Quantitative CLGB measures extracted for three separate imaging cohorts.

| | High Resolution 760 $\mu\text{m}$<br>(average of single subject<br>across 3 sessions) | | | HCP Test Retest<br>(average across subjects<br>first scan only, N=44) | | | Clinical Test Retest<br>(average across subjects<br>first scan only, N=24) | | |
| --- | --- | --- | --- | --- | --- | --- | --- | --- | --- |
|  | Left | Right | p-value | Left | Right | p-value | Left | Right | p-value |
| Volume<br>($\text{mm}^3$) | 2841 ±<br>223 | 4031 ±<br>111 | 0.009 | 3443 ±<br>416 | 4388 ±<br>523 | < 0.001 | 5026 ±<br>721 | 6161 ±<br>886 | < 0.001 |
| FA | 0.43 ±<br>0.006 | 0.45 ±<br>0.003 | 0.04 | 0.41 ±<br>0.025 | 0.39 ±<br>0.018 | < 0.001 | 0.33 ±<br>0.020 | 0.35 ±<br>0.020 | < 0.001 |
| fODF lobe<br>integral | 0.46 ±<br>0.002 | 0.39 ±<br>0.004 | 0.014 | 0.26 ±<br>0.020 | 0.30 ±<br>0.024 | < 0.001 | 0.29 ±<br>0.022 | 0.36 ±<br>0.029 | < 0.001 |
| Weighted<br>FA | 0.024 ±<br>0.004 | 0.028 ±<br>0.003 | **0.32** | 0.07 ±<br>0.013 | 0.05 ±<br>0.010 | < 0.001 | 0.05 ±<br>0.010 | 0.04 ±<br>0.009 | 0.014 |
| Weighted<br>fODF lobe<br>integral | 0.023 ±<br>0.003 | 0.021 ±<br>0.003 | **0.38** | 0.033 ±<br>0.006 | 0.031 ±<br>0.005 | 0.016 | 0.030 ±<br>0.007 | 0.033 ±<br>0.008 | **0.083** |

### 3.5 Comparison to alternative CLGB targeted tractography approaches

Two alternative tractography approaches targeting CLGBs were compared to the proposed surface-based seeding method, namely surface-based seeding along the striatum but with a random initial direction (No Normal) and seeding in a random direction/location within an internal capsule ROI. An example CLGB reconstruction for the left striatum of a single subject is visualized in Figure 10 for each method. Compared to the other approaches, the proposed method (Surface + Normal) generates more dense CLGB-streamline coverage throughout the internal capsule. In addition, the proposed method successfully reconstructs CLGB-streamlines along the posterior end of the striatum, a region where the alternative approaches fail to reconstruct any streamlines. Even though seeding from the surface without the normal produced a similar number of streamlines compared to the proposed method, most of these streamlines were oriented in the inferior superior direction along the walls of the internal capsule and were not representative of CLGBs. Streamline count, average streamline density (within the internal capsule ROI) and voxel coverage (within the internal capsule ROI) are reported in Table 3. Similar to qualitative observations, the proposed method generated more CLGB-streamlines and had a higher streamline density compared to the alternative approaches (p < 0.001). Voxel coverage was higher for the proposed method compared to seeding randomly from the internal capsule ROI (p < 0.001). Whereas voxel coverage was lower for the proposed method compared to seeding directly from the surface with no direction (p < 0.001), again reflecting the large amount of inferior-superior non-CLGB streamlines reconstructed when no initial seeding direction is provided.

**Figure 10.**
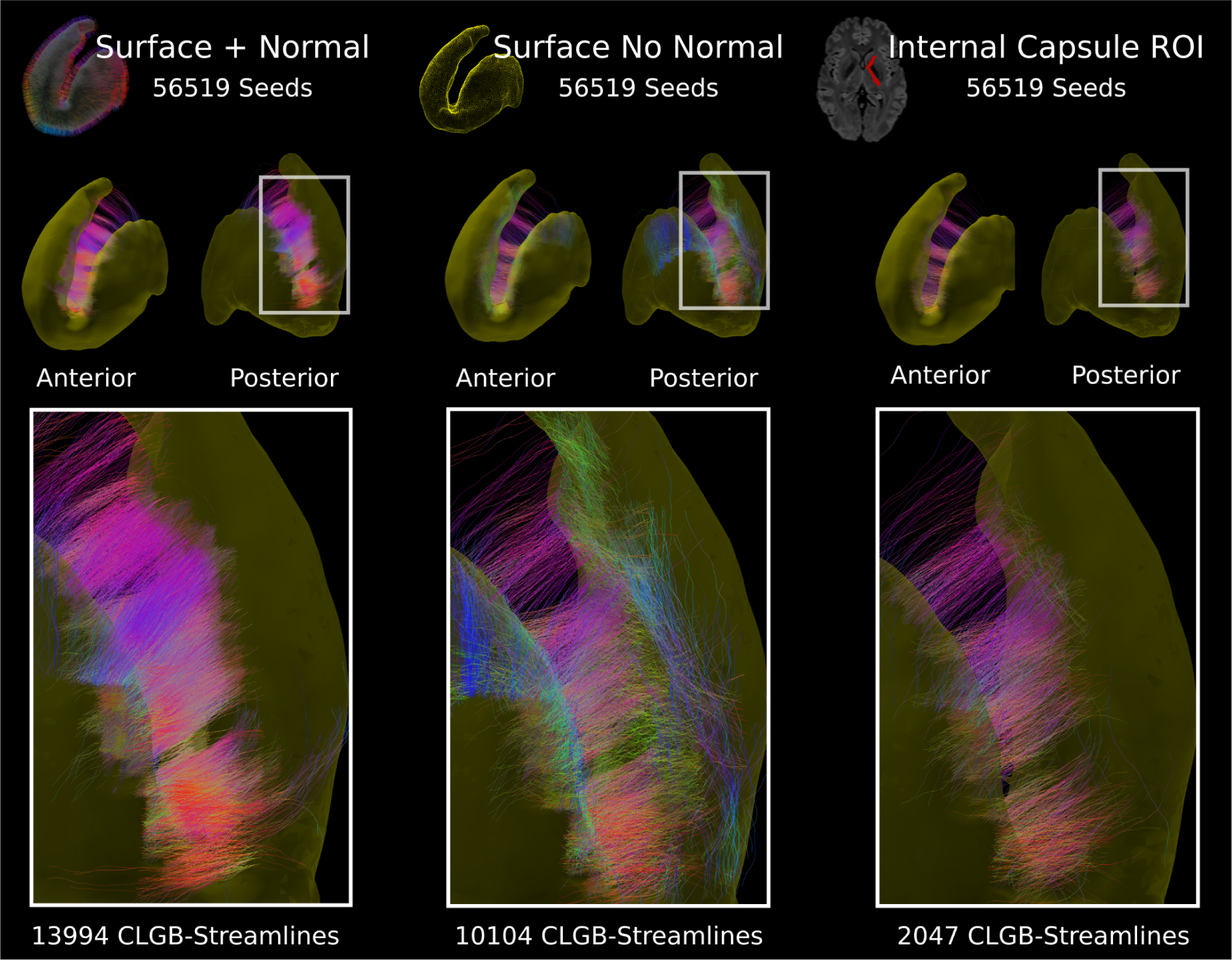
Comparison between the proposed CLGB reconstruction method (Surface + Normal) and two alternative approaches, seeding directly from the surface with a random initial direction (Surface No Normal) and seeding in a random direction/location within an internal capsule region of interest (ROI). The proposed method produced more dense CLGB-streamline coverage throughout the internal capsule compared to the other two methods and was able to reconstruct thinner CLGBs at the posterior end of the striatum. Both of the alternative methods produced a smaller amount of CLGB-streamlines compared to the proposed method and seeding directly from the surface without the initial direction produced a large amount of streamlines oriented in the inferior superior direction not representative of CLGBs.

**Table 3.**
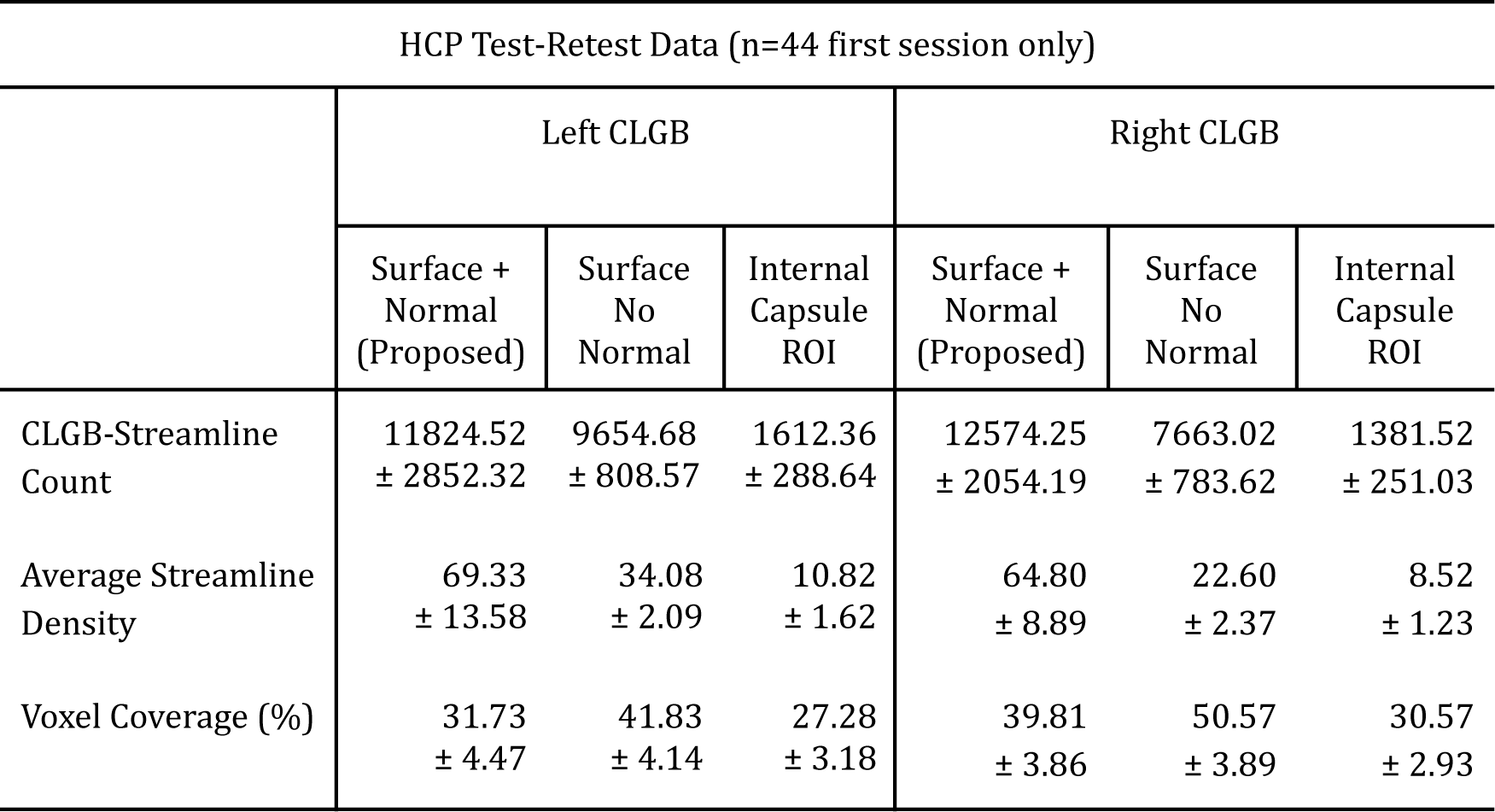
Tractography streamline test-retest comparison between the proposed surface-based seeding CLGB reconstruction method and two alternative approaches.

| HCP Test-Retest Data (n=44 first session only) |  |  |  |  |  |  |
| --- | --- | --- | --- | --- | --- | --- |
|  | Left CLGB |  |  | Right CLGB |  |  |
|  | Surface +<br>Normal<br>(Proposed) | Surface<br>No<br>Normal | Internal<br>Capsule<br>ROI | Surface +<br>Normal<br>(Proposed) | Surface<br>No<br>Normal | Internal<br>Capsule<br>ROI |
| CLGB-Streamline<br>Count | 11824.52<br>± 2852.32 | 9654.68<br>± 808.57 | 1612.36<br>± 288.64 | 12574.25<br>± 2054.19 | 7663.02<br>± 783.62 | 1381.52<br>± 251.03 |
| Average Streamline<br>Density | 69.33<br>± 13.58 | 34.08<br>± 2.09 | 10.82<br>± 1.62 | 64.80<br>± 8.89 | 22.60<br>± 2.37 | 8.52<br>± 1.23 |
| Voxel Coverage (%) | 31.73<br>± 4.47 | 41.83<br>± 4.14 | 27.28<br>± 3.18 | 39.81<br>± 3.86 | 50.57<br>± 3.89 | 30.57<br>± 2.93 |

## 4 Discussion

The CLGBs crossing the internal capsule are associated with an organized anisotropic molecular diffusion signal. This anisotropic signal is oriented orthogonal to the highly anisotropic diffusion signal corresponding to the WM of the internal capsule. When applying a standard CSD local modeling pipeline, “secondary” peaks in the fODF lobes are observed that form a striate pattern in the internal capsule resembling CLGBs. Here a novel tractography seeding strategy is proposed that uses the geometry of the striatum surface to reconstruct CLGBs by seeding orthogonally from the striatum surface to traverse the fODF peaks associated with the CLGBs. Using this method, CLGB-streamline reconstructions clearly overlap the GM of the CLGBs on high resolution 760 µm isotropic diffusion data, demonstrating that the anisotropic diffusion associated with the CLGBs is continuously organized parallel to these structures. The same CLGB reconstruction method can also be applied with high repeatability using more feasible lower resolution diffusion acquisitions (e.g. 1.25 mm and 2 mm isotropic). Overall, this work demonstrates the organized molecular water diffusion profile of the CLGBs and proposes an effective method for reconstructing and quantifying the diffusion signal associated with the CLGBs.

### 4.1 Reconstruction and measurement of CLGBs with diffusion MRI

This is the first work that reports an anisotropic diffusion MRI signal oriented orthogonally across the internal capsule in regions directly associated with the gray matter of the CLGBs. The proposed surface-based tractography method is able to repeatedly reconstruct CLGBs from diffusion data acquired at 760 µm isotropic resolution, with streamlines that overlap regions corresponding to gray matter, indicating that the diffusion anisotropy of the CLGBs persists across the entire structure forming a unique configuration of GM that passes through the WM of the internal capsule. The anisotropy of the diffusion signal in these regions also implies a general microscopic organization of the CLGBs oriented across the internal capsule, which is unique when compared to the fODFs generated for the core of the putamen and caudate nucleus. To further explain a potential source of this unique anisotropic diffusion signal originating in the CLGBs, a publicly available non-human primate immunohistochemistry dataset was investigated (MacBrain Resource: https://medicine.yale.edu/neuroscience/macbrain/). Regions of interest for CLGB, putamen and caudate nucleus regions are visualized on serial coronal sections from a female rhesus monkey (*Macaca mulatta*, 8.07 years old) that were stained for different proteins (Figure 11). From neuronal nuclear protein (Neu) and Nissl-stained sections, compared to the putamen and caudate nucleus, the CLGB region showed similar neuronal density. However, neuronal cell bodies and dendrites stained for (neurofilament protein - SMI-32 and parvalbumin - PV) as well as myelinated axons (stained for myelin basic protein - MBP), demonstrated a coherent pattern of neurites (primarily non-myelinated) organized parallel to the CLGBs. These observations were consistent across multiple specimens (data not shown). Taken together, they suggest that organized dendritic bundles across the CLGBs may explain the anisotropic diffusion signal observed in the current study. Although more work is needed to confirm the major source of the anisotropic diffusion signal in the CLGBs, it should be noted that diffusion anisotropy is not only observed in myelinated tissue (Beaulieu, 2002) and that coherently organized fibers have been speculated previously to be a major source of diffusion anisotropy in the cortical GM (Calamante et al., 2018; McNab et al., 2013).

**Figure 11.**
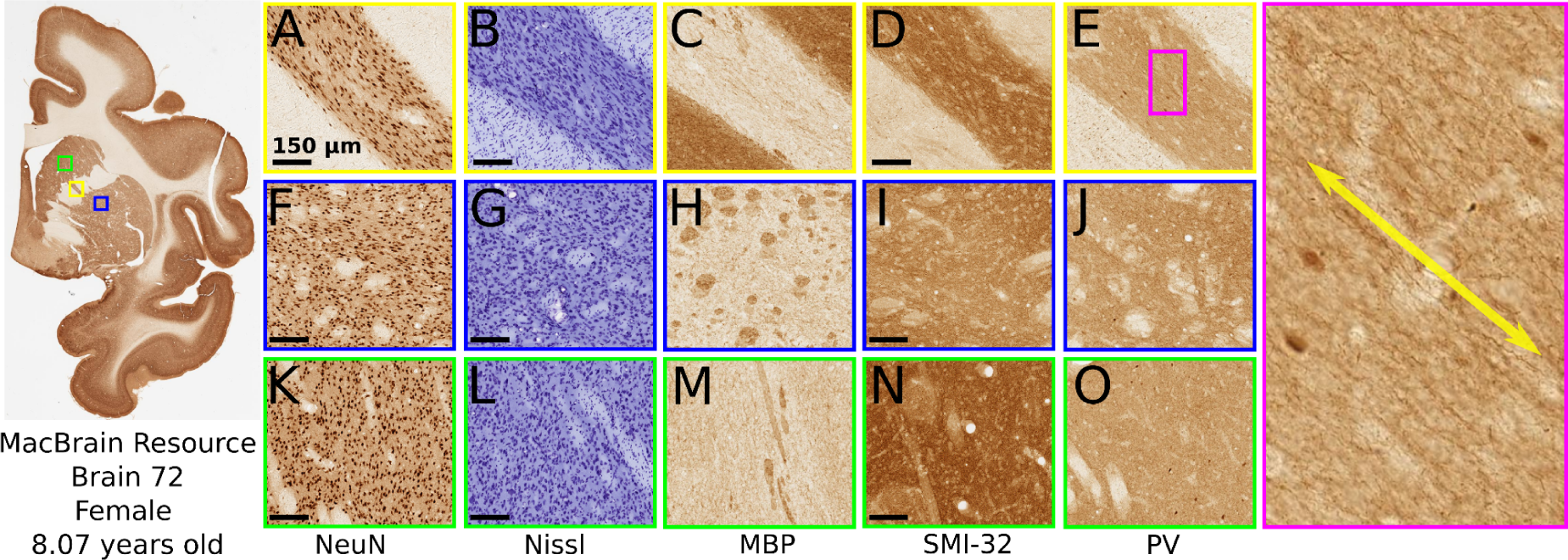
Immunostaining performed on serial coronal sections of a rhesus monkey brain (*Macaca mulatta*, female, 8.07 years old, 14 stainings, 1 mm interval). Higher magnifications are shown for a single CLGB (yellow), a region in the putamen (blue) and a region in the caudate nucleus (green). Staining for neuronal cell bodies (neuronal nuclear protein - NeuN and Nissl) showed similar neuronal density in the CLGB region (A, B) compared to the putamen (F, G) and caudate nucleus (K,L). Immunostaining for both neuronal cell bodies and dendrites (monoclonal antibody to neurofilament protein - SMI-32 and parvalbumin - PV) as well as staining for myelinated axons (myelin basic protein - MBP) show a coherent pattern of fibers oriented parallel (yellow arrow) to the CLGB (D,E zoomed pink region)) a pattern that is shared by a small but coherently oriented group of myelinated axons (C). The coherent fiber pattern was not observed for the control putamen (H, I, J) or caudate nucleus regions (M, N, O).

Given that the CLGBs are likely the only coherently organized structure oriented across the superior portion of the anterior limb of the internal capsule, the diffusion anisotropy associated with the CLGBs is isolated and thus can be detected at both high-resolution as well as at more practically acquirable resolutions. This would suggest that although volumetric properties of the CLGBs may only be measurable at high resolutions smaller than ∼1.0 mm isotropic like performed previously using structural imaging (Dang et al., 2023), the diffusion signal is sensitive to CLGB microstructure at more coarse resolutions. Hence, dMRI could become a useful method for probing the microstructure of these connections in healthy individuals and those with pathology. As a first attempt at measuring the diffusion anisotropy of CLGBs, the direction/density of the CLGB-streamline reconstructions were used to extract a fixel-based measure of the small fODF amplitudes associated with these structures. By using the CLGB-streamline reconstruction to measure the CLGB fODF peaks, it is anticipated that a partial measurement of the much larger fODFs associated with the WM of the internal capsule is reduced. In these measurements of diffusion tensor/fODF parameters the streamline weighted fODF lobe integral measurement did not show a strong hemispheric (left-right) difference compared to the other chosen metrics, reducing the sensitivity to observed volumetric differences between hemispheres. Thus, using the novel tractography method to first identify the orientation of the fODF lobes of interest may become a useful method for extracting a meaningful dMRI measurement of microstructure from the CLGBs.

### 4.2 Technical details of surface-based tractography seeding

By seeding in the direction outwards and normal to the striatum surface, the proposed method reduces the probability of tracking the larger fODF lobes belonging to the WM of the internal capsule and increases the probability of tracking within the smaller fODF lobes associated with the CLGBs. This enabled repeatable streamline reconstructions of the CLGBs across multiple datasets with weighted Dice scores (e.g. ∼0.95) in line with repeatability measures reported for large WM fiber bundles (Rheault et al., 2022). Importantly, other than the initial seed direction/location only one other probabilistic tracking parameter was modified from default settings, namely the relative spherical function threshold was updated to 0.15 (from 0.1) for the high-resolution and HCP test-retest datasets and remained at 0.1 for the clinical datasets. In testing, increasing the threshold allowed for better separation between CLGBs on the high-res diffusion datasets and a threshold of 0.1 was needed on the clinical data to reconstruct CLGB-streamlines through the smaller fODF amplitudes observed on data with a lower signal-to-noise ratio. Based on these observations, this threshold should be selected carefully and might need to be tuned to datasets of different quality.

In the current study, the proposed surface-based seeding method was compared against two other comparable approaches, namely, seeding at the surface interface without an initial orientation and seeding randomly in an ROI of the internal capsule. Keeping all parameters the same, the proposed method was much more efficient (Left: ∼7 times more CLGB-streamlines, right: ∼ 9 times more CLGB-streamlines) than seeding randomly within an internal capsule ROI. Given the lack of “gold standard” in this experiment a true comparison of reconstruction accuracy between methods was not possible, but this experiment does demonstrate the advantage of the proposed surface-based seeding approach compared to the ROI approach that would take 7 times more seeds (i.e. far more computation time) and in doing so generate a larger number of false positive streamlines that would need to be filtered out. The comparison of the proposed method to seeding from the surface without an initial direction demonstrated the clear advantage of using the surface normal to seed tracking. Even though seeding from the mesh without an initial direction generated a similar number of streamlines to the proposed method, the majority of streamlines were oriented inferior/superior and not associated with the desired orientation of the CLGBs. Thus, seeding from the GM/WM interface of the striatum should be approached with caution in a similar fashion to seeding tractography algorithms from the GM/WM interface of the cortex (Schilling et al., 2017; St-Onge et al., 2018).

Outside the scope of reconstructing CLGBs with streamlines tractography, the proposed surface-based seeding technique also might be useful for reconstructing short range WM connections between subcortical GM structures. Future work will aim to assess the use of the proposed method for investigating other subcortical to subcortical connections such as the pallidothalamic tract (Kwon et al., 2021; Rozanski et al., 2017), or to investigate the subcortical GM connectome (Kai et al., 2022). It is anticipated that this approach could also be used to study projection fibers connecting subcortical GM regions and the cortex, where other methods have used subcortical GM surface models to effectively filter streamlines (Marrakchi-Kacem et al., 2013) and create anatomically plausible stopping criteria for tractography algorithms (Yeh et al., 2017).

### 4.3 Study Limitations

Modeling the diffusion signal from non-exchanging cellular/extracellular compartments of WM is a much less complex task than the open problem of modeling GM where molecular exchange exists between compartments (Jelescu et al., 2022). In the current study, no attempt was made to explicitly model the diffusion properties of GM or use a GM response function during CSD. Instead, a classical WM modeling pipeline was used which relied on a single WM response function and measured the geometry of the resulting fODF lobes in the direction of the CLGB-streamline reconstructions. In testing, multi-tissue CSD (Jeurissen et al., 2014) was performed and classified voxels containing CLGBs as WM, likely because this modeling technique treats GM as an isotropically diffusing compartment unfit for the anisotropic diffusion signal of the CLGBs (data not shown). Compared to the CSD based approach, other modeling techniques that do not require the strong assumption of a signal response function might be beneficial for accurately characterizing the diffusion signal within the CLGBs (e.g. model-free mean apparent propagator (MAP) MRI (Özarslan et al., 2013)). Outside of the scope of the current study, future work will aim to accurately model the anisotropic GM of the CLGBs within the internal capsule. That being said, although a microstructural interpretation of the presented fODF lobe based method is limited, it is anticipated that this measure might be sensitive to thinning or pruning of the CLGBs in aging or in neurodegenerative conditions where the caudate nucleus and putamen may be implicated.

Here, a custom segmentation pipeline to segment the striatum and create a WM mask was used. This ensured that tractography seeds generated from the proposed surface-based method were well aligned to the WM/GM interface on the diffusion images. Other surface-based segmentation methods such as FSL FIRST (Patenaude et al., 2011), would likely also be adequate as input for generating the required tractography seeds, however given segmentation methods are usually designed to work on whole brain T1-weighted images an accurate registration would be a prerequisite for their use with the proposed approach. Importantly, standard subcortical GM segmentation methods including the one used in the current study exclude a large portion of the tail of the caudate nucleus which extends posteriorly and curves downward eventually running superior to the hippocampus and terminating in the amygdala. On high resolution (100 micron isotropic) T2 weighted imaging, the caudate tail is also clearly connected to the putamen via CLGBs that are missed by the proposed tractography methodology a result of the current limitations in subcortical GM segmentation techniques.

## 5 Conclusions

The propagation of water molecules estimated with diffusion MRI is sensitive to the anisotropic organization of the GM tissue constituting the CLGBs crossing the internal capsule. As demonstrated in this study, such anisotropy can be detected at imaging resolutions from 760 µm to 2 mm isotropic using standard CSD fODF modeling. The proposed surface-based tractography seeding method presents a novel technique for reconstructing the CLGBs and extracting a microstructural measurement from the corresponding fODF lobes. Taken together, diffusion MRI combined with surface-based tractography seeding could become a useful technique for understanding the microstructure of CLGBs. Given this technique can be applied at more feasible imaging resolutions typically acquired in clinical research, it has the potential to be applied in future in-vivo studies targeting clinical pathologies, aging or neurodegeneration.

## Data Code and Availability

The software for the novel tractography seeding method will be made available upon publication. The locally acquired diffusion dataset used in the current study are not publicly available because the data is not exclusively owned by the authors. However, given a reasonable request to the last author they are available from the corresponding under data sharing agreements with the owners. All other data used in the current study are publicly available from the referenced resources.

## Author Contributions

**Graham Little:** Writing - original draft, Conceptualization, Methodology, Data curation, Formal analysis, Software. **Charles Poirier:** Software, Writing - review & editing **Martin Parent:** Methodology, Writing - review & editing. **Laurent Petit:** Conceptualization, Methodology, Writing - review & editing. **Maxime Descoteaux:** Conceptualization, Methodology, Writing - original draft, Writing - review & editing.

## Acknowledgements

The authors would like to thank the following institutions for funding portions of this work; the Natural Sciences and Engineering Research Council of Canada (salary GL and CP), Unifying Neuroscience and Artificial Intelligence - Québec (GL), Centre de recherche du Centre hospitalier universitaire de Sherbrooke (GL), Fonds de Recherche du Québec Nature et technologies (CP) and the Université de Sherbrooke (CP). Part of this research was supported by the NSERC Discovery grant and the Université de Sherbrooke Institutional Chair in Neuroinformatics from Pr Descoteaux. The funders had no role in study design, data collection, analysis, decision to publish, or preparation of the manuscript.

